# Use of multivariable Mendelian randomization to address biases due to competing risk before recruitment

**DOI:** 10.1101/716621

**Authors:** C Mary Schooling, Priscilla M Lopez, Zhao Yang, J V Zhao, SL Au Yeung, Jian V Huang

## Abstract

**Background:** Mendelian randomization (MR) provides unconfounded estimates. MR is open to selection bias particularly when the underlying sample is selected on surviving the genetically instrumented exposure and other conditions that share etiology with the outcome (competing risk before recruitment). Few methods to address this bias exist.

**Methods:** We use directed acyclic graphs to show this selection bias can be addressed by adjusting for common causes of survival and outcome. We use multivariable MR to obtain a corrected MR estimate, specifically, the effect of statin use on ischemic stroke, because statins affect survival and stroke typically occurs later in life than ischemic heart disease so is open to competing risk.

**Results:** In univariable MR the genetically instrumented effect of statin use on ischemic stroke was in a harmful direction in MEGASTROKE and the UK Biobank (odds ratio (OR) 1.33, 95% confidence interval (CI) 0.80 to 2.20). In multivariable MR adjusted for major causes of survival and ischemic stroke, (blood pressure, body mass index and smoking initiation) the effect of statin use on stroke in the UK Biobank was as expected (OR 0.81, 95% CI 0.68 to 0.98) with a Q-statistic indicating absence of genetic pleiotropy or selection bias, but not in MEGASTROKE.

**Conclusion:** MR studies concerning late onset chronic conditions with shared etiology based on samples recruited in later life need to be conceptualized within a mechanistic understanding, so as to any identify potential bias due to competing risk before recruitment, and to inform the analysis and interpretation.

## Introduction

Mendelian Randomization (MR), i.e., instrumental variable analysis with genetic instruments, is an increasingly popular and influential analytic technique [1, 2], which can be used to investigate causal effects even when no study including both exposure and outcome of interest exists. Invaluably, MR studies have provided estimates more consistent with results from randomized controlled trials (RCTs) than conventional observational studies, even foreshadowing the results of major trials [3]. MR studies are often presented as observational studies analogous to RCTs [4, 5] because they take advantage of the random assortment of genetic material at conception, while observational studies are open to biases from confounding and selection bias [6]. Instrumental variable analysis is described in health research as addressing confounding [7, 8], i.e., bias from common causes of exposure and outcome [6]. MR is currently described as “less likely to be affected by confounding or reverse causation than conventional observational studies” [2].

MR was originally thought to be less open to selection bias than conventional observation studies [9]. Selection bias is now increasingly widely recognized as a limitation of MR [10-20] which may violate the instrumental variable assumptions. Sources of potential selection bias in MR have been specifically identified as selecting an unrepresentative sample [10, 12, 13], attrition from an initially representative sample, such as a birth cohort [13], and selecting a sample strongly on surviving the exposure [11] or genotype of interest [16, 21]. What has not explicitly been considered is selecting the underlying sample(s) on surviving the genotype of interest in the presence of competing risk of the outcome. MR studies are particularly vulnerable to sample selection on survival because of the time lag between genetic randomization (at conception) and typical recruitment into genetic studies of major diseases in middle- to old-age. MR studies also often concern major causes of death thought to share considerable etiology. For example, lipids, blood pressure, diabetes, lifestyle (such as smoking, diet, physical activity and sleep) and socio-economic position cause both ischemic heart disease (IHD) and ischemic stroke, with death from IHD typically occurring at younger ages than death from stroke [22, 23]. As a result, a study of the association of lipid modifiers with stroke among the living will automatically select on surviving high lipids and on surviving competing risk of prior death from IHD due to shared etiology between IHD and stroke. Some people dying from genetically high lipids and others dying from IHD before recruitment into a stroke study will leave a shortage of people available to recruit with genetically high lipids and susceptibility to stroke, thereby obscuring any effect of lipids or lipid modifiers on stroke. Correspondingly, MR studies suggest less effect of lipids and lipid modifiers on stroke than IHD [24, 25], although RCTs suggest similar effects [26-28]. Similarly, MR studies do not consistently show detrimental effects of body mass index (BMI) on stroke [29]. In this study, we explain how potential violations of the instrumental variable assumptions due to inadvertently recruiting survivors of the exposure and competing risk of the outcome may bias MR estimates. We explain how to correct for this bias using multivariable MR and provide a simple means of estimating how large the bias is likely to be.

## Methods

### Potential biasing pathways due to recruiting on selective survival

Figure 1a shows the directed acyclic graph for MR illustrating the instrumental variable assumptions typically referred to as relevance, independence and exclusion-restriction. Relevance is explicitly indicated by the arrow from instrument to exposure. Independence is implicitly indicated by the lack of an arrow from confounders of exposure on outcome to instrument. Exclusion-restriction is implicitly indicated by the lack of arrows linking instrument to outcome, sometimes illustrated as no arrow from instrument to outcome indicating no pleiotropy [30-33] (Figure 1b). Figure 1c shows selection on survival of both instrument and common causes of the outcome (U_2_) [12, 19], which also violates the exclusion restriction assumption, particularly when stated as “*every unblocked path connecting instrument and outcome must contain an arrow pointing into the exposure*” [34]. Figure 1d explicitly shows survival on instrument, and another disease (Y_2_) sharing etiology (U_2_) with the outcome (Y_1_). Figure 1e shows the exclusion restriction assumption with both no pleiotropy and no selection bias from competing risk made explicit. Notably, Figures 1c, d and e are very similar in structure to a well-known example of selection bias which occurs when conditioning on an intermediate (or covariable adjustment) reverses the direction of effect: the “birth weight” paradox [35]. In the birth weight paradox adjusting the association of maternal smoking with infant death makes maternal smoking look protective; further adjusting for all common causes of birth weight and infant survival, thought to be birth defects, should remove this bias [35] by blocking the path from survival to outcome. Similarly, bias due to inadvertently selecting the underlying sample in an MR study on surviving the genetically instrumented exposure and surviving competing risk of the outcome should be ameliorated by adjusting for major causes of survival and the outcome (Figure 2). The recent development of multivariable MR [36] provides the means to do so. Specifically, as indicated in Figure 1(c-d), where the univariable MR may be biased using multivariable MR adjusting for the main determinants of survival and outcome may reduce bias by at least partially blocking any backdoor paths from instrument to outcome.

**Figure 1:**
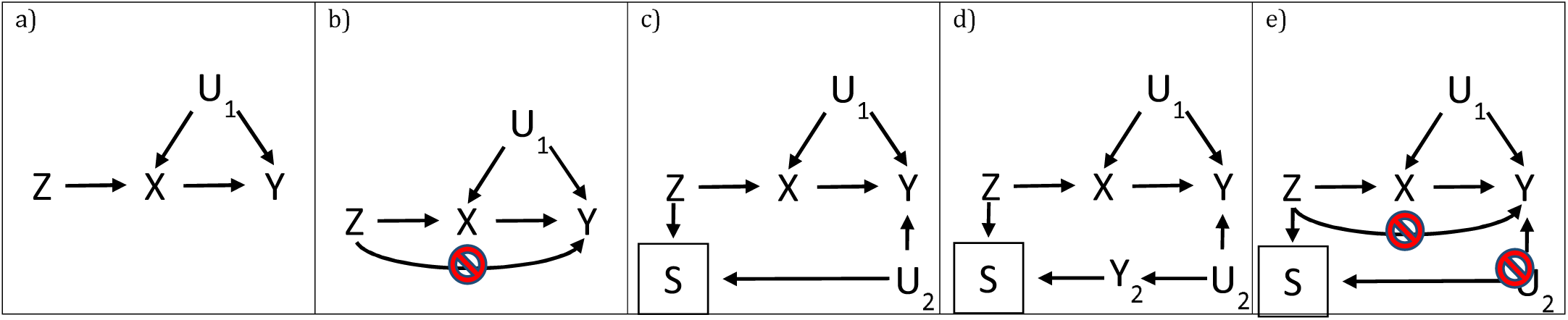
Directed acyclic graphs with instrument (Z), outcome (Y), exposure (X), confounders (U) and survival (S), where a box indicates selection, for a) a valid Mendelian randomization study, and b) a Mendelian randomization study with an invalid instrument through violation of the exclusion-restriction assumption via pleiotropy, c) a Mendelian randomization study with an invalid instrument through violation of the exclusion-restriction assumption via survival on instrument and shared etiology with the outcome (U_2_), d) a Mendelian randomization study with an invalid instrument through violation of the exclusion restriction assumption via survival (S), competing risk of another disease (Y_2_) and shared causes (U_2_) with (Y_2_) and the outcome (Y) and e) a Mendelian randomization illustrating both conditions which have to be met to satisfy the exclusion restriction assumption.

**Figure 2:**
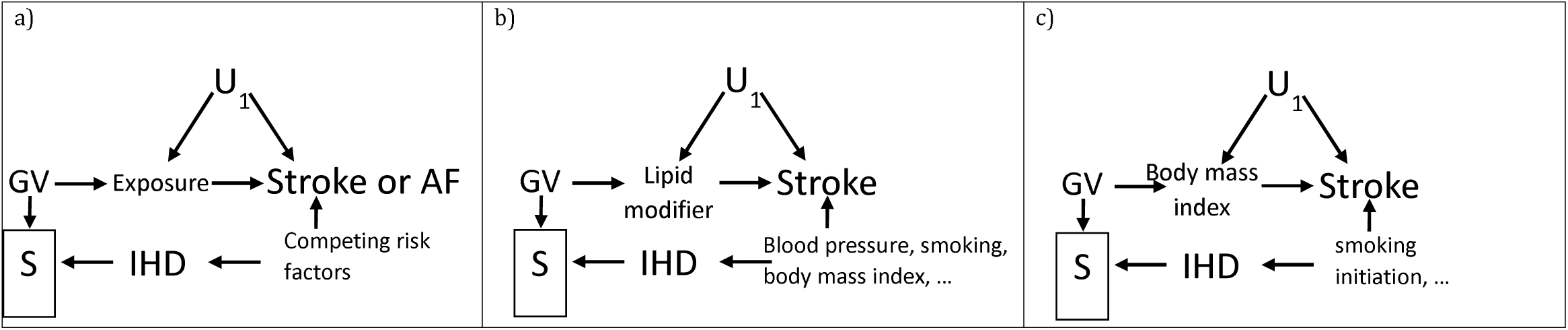
Directed acyclic graphs showing how selection bias could occur because of selection on survival (S), indicated by a box, on the instrument (GV) and on competing risk of ischemic heart disease (IHD) which shares causes with the outcome of interest, i.e., stroke, with U_1_ as confounders of exposure and outcome, when assessing a) effects of an exposure on stroke or AF, b) effects of lipid modifiers on stroke and b) effects of body mass index on stroke.

In addition, to provide triangulation, the level of selection bias due to surviving to recruitment on genetically instrumented exposure in the presence of competing risk of the outcome can also be thought of as depending on the proportion of the exposed who are not available for recruitment because of prior death due to the genetically predicted exposure and the proportion of those who could have experienced the outcome who are not available for recruitment because of prior death from a competing risk. Assuming these proportions are independent and their corresponding probabilities do not sum to more than 1, then for an observed odds ratio greater than 1 the true odds ratio for genetically predicted exposure on disease can be estimated as the observed odds ratio multiplied by the ratio of the probability of surviving the exposure and the competing risk to the probability of surviving the exposure or the competing risk, as shown in the Appendix.

### Examples of selection bias and amelioration

We investigated effects of lipid modifiers and BMI on ischemic stroke (IS) as possible exemplars, because previous MR studies of these exposures on stroke have not always given the expected results [29, 37]. Statins and PCSK9 inhibitors are very well-established interventions for cardiovascular disease, which reduce low density lipoprotein (LDL)-cholesterol, IHD [26-28], stroke [26-28] and atrial fibrillation (AF) [38]. BMI is also known to be harmful. IHD, IS and AF also share major causes independent of lipid modifiers, such as blood pressure [39, 40], smoking, lifestyle and socio-economic position. Death from IHD typically occurs at earlier ages than death from IS in Western populations [22, 23]. AF may also be a consequence of IHD. Figure 2 suggests bias would be expected for harmful exposures on IS or AF. Adjusting for major factors causing survival to recruitment into the underlying studies of IS or AF, as shown for lipid modifiers on IS (Figure 2b) or BMI on IS (Figure 2c) should reduce the bias. As such, univariable MR, even with well-defined genetic instruments free from genetic pleiotropy, might generate biased estimates due to selection bias violating the exclusion-restriction assumption, but appropriate use of multivariable MR might ameliorate the problem.

We used well-established independent genetic variants for statins (rs12916) and PCSK9 inhibitors (rs11206510, rs2149041 and rs7552841) [41], and BMI (96 variants) [42]. Using two-sample univariable MR we applied these variants to major GWAS in people largely of European descent of IHD (CARDIoGRAMplusC4D 1000 Genomes) [43], IS (MEGASTROKE) [44] and AF (Nielsen et al) [45]. We also used the UK Biobank summary statistics for IHD and IS [46], but not for AF because the AF GWAS includes the UK Biobank data [45]. We obtained univariable MR estimates by meta-analyzing the Wald estimates (genetic variant on outcome divided by genetic variant on exposure) using inverse variance weighting, with multiplicative random effects, after aligning variant estimates on the same effect allele in each study.

We used multivariable two-sample MR to adjust lipid modifiers on IS and AF for major causes of survival (smoking initiation [47], blood pressure and BMI)[48, 49] and IS, and to adjust BMI on IS for smoking initiation. We used published independent genetic instruments for smoking initiation (327 variants) [47], systolic (SBP) and diastolic blood pressure (DBP) (novel replicated variants (SBP 130, DBP 91)) [50] and BMI (96 variants) [42]. Genetic associations for all the instruments with low-density cholesterol, ever smoking, SBP, DBP, and BMI were obtained from the UK Biobank summary statistics (http://www.nealelab.is/uk-biobank) adjusted for age, sex, age^2^, sex*age, sex* age^2^ and the first 20 principal components. We used the MR-Base ld_clump R package with r^2^<0.05 to obtain independent genetic variants and the Mendelianrandomization package to obtain IVW multivariable estimates. We reported the multivariable conditional F-statistic as a measure of instrument strength and the multivariable Q-statistic as a measure of instrument pleiotropy [36], obtained using the MVMR package [36]. Notably, in this context a significant multivariable Q statistic may indicate genetic pleiotropy or violation of the exclusion restriction assumption by selection bias.

This study only used publicly available genetic summary statistics, collected with consent, and so does not require ethical approval.

## Results

As expected, the cases recruited into the underlying GWAS in the main analysis [43-45] seemed to be youngest for IHD and oldest for AF with IS somewhere in between (Supplementary Table 1). In univariable MR genetically instrumented statin or PCSK9 inhibitor use reduced IHD, while genetically instrumented BMI increased IHD (Table 1). Estimates were similar using CARDIoGRAMplusC4D 1000 Genomes and the UK Biobank. IHD is not expected to be majorly open to competing risk, so was not considered further. In univariable MR, genetically instrumented statin or PCSK9 inhibitor use was not associated with a lower risk of IS or AF; some estimates for statins were in the direction opposite to expected (Table 1). In univariable MR, genetically instrumented BMI did not consistently increase IS, but did increase AF (Table 1)

**Table 1:** Effect of genetically instrumented statin and PCSK9 inhibitor [41] (in effect size of LDL-cholesterol) use and BMI [42] on IHD using the CARDIoGRAMplusC4D 1000 Genomes based GWAS [43] and the UK Biobank, on all ischemic stroke using MEGASTROKE [44] and the UK Biobank and on AF using a study by Nielsen et al [45] from univariable Mendelian randomization and from multivariable Mendelian randomization, with genetically predicted statins and PCSK9 inhibitors adjusted for systolic blood pressure [50], diastolic blood pressure [50], smoking initiation [47] and BMI [42], and BMI adjusted for smoking initiation.

| Disease | Source of genetic associations with disease | Exposure | Univariable |  | Multivariable |  | Conditional F | Q p-value |
| --- | --- | --- | --- | --- | --- | --- | --- | --- |
|  |  |  | OR | 95% CI | OR | 95% CI |  |  |
| Ischemic heart disease | CARDIoGRAMplusC4D 1000 Genomes | statin | 0.56 | 0.41 to 0.75 |  |  |  |  |
|  |  | PCSK9 inhibitor | 0.32 | 0.22 to 0.46 |  |  |  |  |
|  |  | BMI | 1.57 | 1.36 to 1.81 |  |  |  |  |
|  | UK Biobank (SAIGE) | statin | 0.69 | 0.52 to 0.93 | 0.64 | 0.55 to 0.75 | 6.7 | 4.5e-23 |
|  |  | PCSK9 inhibitor | 0.47 | 0.34 to 0.65 | 0.63 | 0.54 to 0.73 | 6.8 | 4.6e-23 |
|  |  | BMI | 1.38 | 1.18 to 1.61 | 1.48 | 1.33 to 1.65 | 17.3 | 2.5e-23 |
| All ischemic stroke | MEGASTROKE | statin | 1.18 | 0.84 to 1.65 | 1.05 | 0.90 to 1.23 | 6.6 | 2.3e-8 |
|  |  | PCSK9 inhibitor | 0.94 | 0.65 to 1.37 | 1.05 | 0.90 to 1.21 | 6.8 | 1.7e-8 |
|  |  | BMI | 1.18 | 1.04 to 1.34 | 1.16 | 1.05 to 1.28 | 17.2 | 0.001 |
|  | UK Biobank (SAIGE) | statin | 1.33 | 0.80 to 2.20 | 0.77 | 0.63 to 0.95 | 6.7 | 0.18 |
|  |  | PCSK9 inhibitor | 0.96 | 0.55 to 1.69 | 0.77 | 0.62 to 0.94 | 6.8 | 0.20 |
|  |  | BMI | 1.13 | 0.93 to 1.36 | 1.27 | 1.10 to 1.47 | 17.3 | 0.02 |
| Atrial fibrillation | Nielsen et al (including UK Biobank) | statin | 1.22 | 0.97 to 1.54 | 1.14 | 1.01 to 1.29 | 6.7 | 1.3e-39 |
|  |  | PCSK9 inhibitor | 0.79 | 0.62 to 1.01 | 1.09 | 0.96 to 1.23 | 6.8 | 2.7e-40 |
|  |  | BMI | 1.46 | 1.34 to 1.59 | 1.44 | 1.34 to 1.56 | 17.2 | 1.5e-15 |

In multivariable MR, the conditional F-statistics for each exposure were similar in each analysis, suggesting similar instrument strength (Table 1). The Q-statistics were not significant for lipid modifiers or BMI on UK Biobank stroke (Table 1), indicating these estimates were unlikely open to bias from genetic pleiotropy or from selection bias. The multivariable MR estimates in the UK Biobank, in contrast to the corresponding univariable MR estimates, showed that genetically instrumented lipids modifiers protected against stroke and that genetically instrumented BMI caused stroke (Table 1). The Q-statistics were highly significant for lipid modifiers and BMI on MEGASTROKE IS and AF (Table 1), indicating these estimates were likely still biased by pleiotropy probably from selection bias given the same instruments gave apparently unbiased for stroke in the UK Biobank. Correspondingly, the multivariable estimates were similar to the univariable estimates, and for lipid modifiers differed from those expected from RCTs (Table 1).

To provide triangulation we estimated whether the level of selection bias for statins on IS, from surviving genetically instrument statins and IHD, was consistent with the univariable estimate. The odds ratio for the protective allele of the statin SNP (rs12916) on IHD used to obtain the Wald estimate was 0.96. Assuming statins have the same effect on IHD and IS, it would only take 10% with that harmful allele and 25% of potential IS cases to have died from IHD or other competing risk before recruitment into a stroke study for the observed odds ratio to be exactly 1.0, which would give a null MR estimate. If instead 40% of potential IS cases had died from competing risk before recruitment then the odds ratio would reverse to 1.04, and give an MR estimate similar to the univariable estimate from MEGASTROKE.

## Discussion

Here, we have shown theoretically, empirically and analytically that univariable MR studies can be open to quite severe selection bias likely arising from selective survival on genetically instrumented exposure when other causes of survival and outcome exist, i.e. competing risk before recruitment. We have also explained the relevance of this situation to the assumptions of MR, as a violation of the exclusion restriction assumption, how to mitigate this bias using multivariable MR, how to assess the success of this mitigation (using the multivariable Q statistic), and how to make an assessment of the possible level of bias using an approximation based on contextual knowledge. Notably, genetic studies are particularly vulnerable to bias because most genetic estimates are of small magnitude; the closer the true estimate is to the null the easier it is for a reversal to occur (Appendix Figure 1).

Our study differs from many other studies suggesting that MR is open to selection bias by specifically identifying when such bias can occur in the context of a typical MR study using existing GWAS, and by showing how any such bias may be addressed along with a means of checking whether the bias has been successfully addressed. For participants selected on surviving the genetically instrumented exposure and competing risk of the outcome, our study is similar to other studies about bias in MR in showing that bias can occur from using GWAS summary statistics with “covariable adjustment” [51]. We add by explaining that selecting from the living is common in MR studies and may engender covariable adjustment on survival. Rather than suggesting that such situations should be avoided [51], precluding MR studies of a harmful exposure on a late-onset disease subject to competing risk, we show how such situations can be addressed. Specifically, external knowledge can be used to identify potential common causes of survival and outcome, followed by multivariable MR to adjust for them and thereby possibly obtain a less biased estimate, bearing in mind the Q statistic. We also show that when it is not possible to adjust for factors causing survival and the outcome the level of potential bias can be estimated. Alternatively, restricting MR studies to younger people will usually reduce bias because death prior to recruitment is less common in younger people. However, these studies may need to consider competing risk after recruitment. Our study also implies that care should be taken in interpreting phenome-wide association studies identifying the effect of a specific genetically instrumented exposure across the phenome, because the effects of harmful exposures observed will vary depending of the level of competing risk of the outcome.

Despite the strengths of our study in explicating and providing means of addressing a relatively common bias in univariable MR, use of multivariable MR requires knowledge of the underlying causal structure and suitable genetic instruments for all sources of bias. Alternative methods to recover from selection bias due to surviving the genetically instrumented exposure and competing risk of the outcome that do not require knowledge of the underlying causal structure or additional data would be easier to use. Second, our study did not conduct simulations of the level of bias. However, such simulations have already been done [51]. The key issue we address is appreciating which real life situations will result in the simulated bias, and what to do to ameliorate it. We provide a means of addressing any such bias using multivariable MR (adjusting for common causes of survival and outcome) as well as a means of assessing the likely validity of the revised estimate (non-significant multivariable Q-statistic). However, as with any bias correction by adjustment, it may not be feasible to recover the correct estimate, due to lack of contextual knowledge, a highly inter-related causal structure, such as the genetic instruments causing common causes of survival and outcome, or a lack of relevant information. So, we also provide a simple analytic means of estimating the level of bias (Appendix 1). Third, we do not provide an exhaustive list of examples of when this bias has occurred, because few MR studies have been validated against RCTs. For example, Alzheimer’s disease usually occurs in old age, and appears to share causes with determinants of longevity [52], so MR studies of Alzheimer’s disease could be open to selection bias but the true causes of Alzheimer’s disease are unknown making any determination of whether the MR studies are biased or not difficult. Finally, the issue of obtaining valid estimates in the presence of selective survival on exposure and competing risk of the outcome is similar to the issue of obtaining valid genetic estimates in other studies of survivors, i.e., patients. The current solution for obtaining valid estimates in genetic studies of patients relies on the assumption that the factors causing disease and disease progression differ [53]. Whether using multivariable MR to adjust observational studies in patients suitably might bear consideration.

## Conclusion

Here, we have shown theoretically, empirically and analytically that univariable MR studies can be open to quite severe selection bias arising from selecting on survival of genetically instrumented exposure when other causes of survival and outcome exist, i.e. competing risk before recruitment. Bias from such selection bias is likely to be least for MR studies of harmless exposures recruited shortly after genetic randomization with no competing risk, i.e., studies using birth cohorts with minimal attrition. Conversely, such bias is likely to be most evident for MR studies recruited at older ages examining the effect of a harmful exposure on an outcome subject to competing risk from shared etiology with other common conditions that occur earlier in life. Use of multivariable MR to adjust for major causes of survival and outcome may ameliorate this bias, while simple sensitivity analysis based on information about the exposure and the natural history of disease may help quantify the magnitude of the bias. Infallible, methods of obtaining valid MR estimates, when the exclusion restriction is invalidated by selection bias stemming from competing risk, that do not require external knowledge, would be helpful.

## Appendix 1: A possible solution for recovering the causal effect in the presence of selection bias due to selecting on surviving the exposure and competing risk of the exposure in a case-control study

The fundamental issue of the selection bias in a case-control study is unknown information for the “missing” (or unselected) participants. **Table 1** shows the possible mechanism generating a biased causal effect due to selection on surviving the exposure (E) and surviving competing risk (CR) of the outcome (D) in a case-control study. Based on the observed data a^’^, b^’^, c^’^, and d^’^, the observed causal effect of E on D using an odds ratio (*0R*^*ObS*^) is,

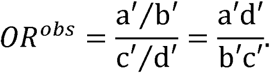

To obtain the true causal effect, we have to recover the data for the whole population, i.e. the birth cohorts who formed the population. Let P_E_ denote the proportion of participants unselected due to E, and let P_CR_ denote the proportion of participants unselected due to CR. Suppose P_E_ and P_CR_ are additive, and 0 < P_E_ + P_CR_ < 1. We can construct the pattern of the unselected participants, as shown in **Table 1**. As such, the causal effect of E on D for the whole population can be estimated as follows,

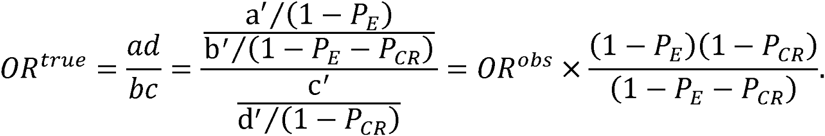

This relationship will be invalid if we replace the OR with a risk ratio.

**Appendix Table 1.** Possible mechanism for biased causal effects in a case-control study due to selection bias from surviving the exposure and competing risk of the outcome

|  |  | S = 1 |  | S = 0 |  |
| --- | --- | --- | --- | --- | --- |
|  |  | D = 1 | D = 0 | D = 1 | D = 0 |
| E = 1 | CR = 1 | a' | b' | P<sub>E</sub> | P<sub>E</sub> + P<sub>CR</sub> |
|  | CR = 0 |  |  |  |  |
| E = 0 | CR = 1 | c' | d' | 0 | P<sub>CR</sub> |
|  | CR = 0 |  |  |  |  |
| Observation |  | OR<sup>obs</sup> |  |  |  |
| Target |  | OR<sup>true</sup> |  |  |  |

Note: S indexes selection status of participants, i.e., S = 1 indicates those selected and S = 0 indicates those unselected. D indexes outcome status, i.e., D = 1 indicates disease and D = 0 indicates no disease. E indexes exposure status, i.e., E = 1 indicates the exposed and E = 0 indicates unexposed. CR indicates competing risk (CR) of the outcome D; i.e., CR = 1 then D = 0, and if D = 1 and CR = 0. a^’^, b^’^, c^’^, and d^’^ are observed data about the selected participants.

Notably, the level of bias depends on the magnitude of the OR. A small OR, of the order of 1.05, as is typical in a genetic study, is much more vulnerable to a reversal of effect from selection bias due to selecting on surviving the exposure and surviving competing risk of the outcome than a larger OR, of the order of 1.50, as is typical in traditional observational studies. To clarify Appendix Figure 1 shows the observed OR plotted against the true OR for different combinations of selection on survival (P_E_) and selection on competing risk of surviving the outcome (P_CR_)

**Appendix Figure 1:**
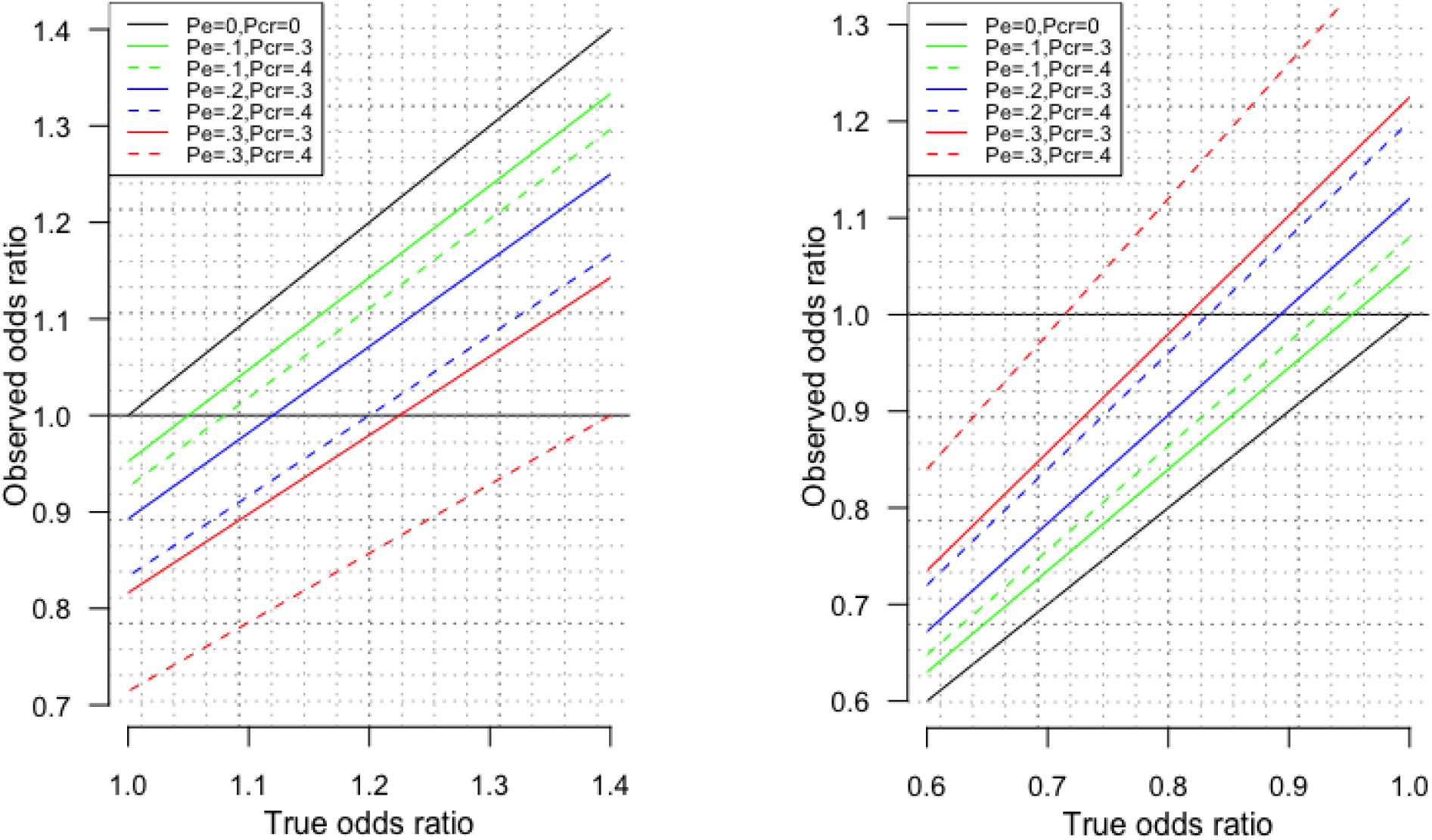
Observed odds ratio against the True odds ratio in the presence of different proportions of death before recruitment due to the exposure (P_E_) and different proportions of death before recruitment due to competing risk of the outcome (P_CR_) for true odds ratios large than 1 (left hand side) and smaller than 1 (right hand side, obtained by taking the inverse of the odds ratio).

**Supplementary table 1:** Study details for the GWAS of IHD, stroke and AF

| Study | Phenotype<br>(Phewas<br>code) | Cases | Non-<br>cases | Mean<br>age of<br>cases | Phenotype definition | Adjusted for (non-<br>genetic) |
| --- | --- | --- | --- | --- | --- | --- |
| Cardiogram<br>1000 genomes<br>GWAS [43] | Ischemic<br>heart disease | 60,801 | 123,504 | n/a,<br>possibly<br>~58 years | “Case status was defined by an inclusive CAD diagnosis (e.g. myocardial infarction (MI), acute coronary syndrome, chronic stable angina, or coronary stenosis >50%)” | Study-specific<br>covariates (not age<br>or sex) |
| UK biobank<br>SAIGE [46] | CAD (411) | 31,355 | 377,103 | n/a | Phewas code based on self-report, hospital episodes and death | Sex, birth year, and<br>principal<br>components 1 to 4 |
|  | Stroke (433) | 8,742 | 399,017 |  |  |  |
|  | AF (427.2) | 14,820 | 380,919 |  |  |  |
| MEGASTROKE<br>[44] | All ischemic<br>stroke | 60,341 | 454,450 | ~69 years | Several different definitions used | Minimum of age<br>and sex |
| AF [45] | Atrial<br>fibrillation | 60,620 | 970,216 | ~74 years | Usually based on ICD-9 427.3 and ICD-10 I48 | Minimum of age<br>(birth year) and sex |

